# Non-invasive real-time access to the output of the spinal cord via a wrist wearable interface

**DOI:** 10.1101/2021.04.06.438640

**Authors:** Irene Mendez, Deren Y. Barsakcioglu, Ivan Vujaklija, Daniel Z. Wetmore, Dario Farina

## Abstract

Despite the promising features of neural interfaces, their trade-off between information transfer and invasiveness has limited translation and viability outside research settings. Here, we present a non-invasive neural interface that provides access to spinal motoneuron activities from a sensor band at the wrist. The interface decodes electric signals present at the tendon endings of the forearm muscles by using a model of signal generation and deconvolution. First, we evaluated the reliability of the interface to detect motoneuron firings, and thereafter we used the decoded neural activity for the prediction of finger movements in offline and real-time conditions. The results showed that motoneuron activity decoded from the wrist accurately predicted individual and combined finger commands and therefore allowed for highly accurate real-time control. These findings demonstrate the feasibility of a wearable, non-invasive, neural interface at the wrist for precise real-time control based on the output of the spinal cord.

## INTRODUCTION

In our ever-growing digital world, human-machine interaction has a pivotal role not only in defining our relationship with technology but also in determining its usability and effectiveness. However, current user input systems, such as keyboards or touchscreens, are constrained by the possibilities of the physical world to capture and deliver users’ intentions. Bypassing these intermediary devices would revolutionize our interaction with technology, making control more intuitive and human centered. Neural interfaces offer a direct decoding of a user’s intentions from the nervous system and thus may enable an intuitive interaction with the environment by translating neural activity into digital inputs to external devices. This approach targets the neural code of motor commands to provide seamless and enhanced control of external systems.

Traditionally, signals from the brain have been leveraged to control prostheses(*1*), wheelchairs(*2*), and exoskeletons(*3*), among other systems(*4, 5*). Nonetheless, direct brain interfaces are currently constrained by a trade-off between their information transfer and invasiveness, with non-invasive systems providing substantially smaller information bandwidth than invasive devices(*6*). Furthermore, acceptance of brain interfaces for daily use outside rehabilitation applications remains uncertain. In comparison, the peripheral nervous system offers a more accessible window to motor volition(*7*). Motoneurons in the spinal cord translate the synaptic inputs they receive from supraspinal centers and peripheral afferents into a neural output sent to the muscles(*8*). The axonal action potentials generated by motoneurons reach the neuromuscular junctions to excite muscle fibers. This generates muscle fiber action potentials that propagate from the neuromuscular junctions to the tendon endings(*9*), generating electric fields detectable on the skin via surface electrodes(*10*). When detected over active muscles, this electrical activity is the electromyographic (EMG) signal, which has been extensively used as input for neuroprostheses(*11*).

The most common approach to human-machine interfacing for decoding distal movements of the upper limb is to record EMG from forearm muscles, yet the least obtrusive and most socially acceptable location for long-term adoption of wearable devices is at the wrist due to social acceptance of wristwatches(*12, 13*). Tendon tissues are dominant at the wrist, and there is minimal muscle mass at the end of several forearm muscles (Fig. 1). Nonetheless, electric fields generated by neural activation can still be detected at the tendons due to volume conduction (Fig. 1). Hereafter, we will refer to signals recorded over tendon tissue as tendon electric signals.

**Fig. 1.**
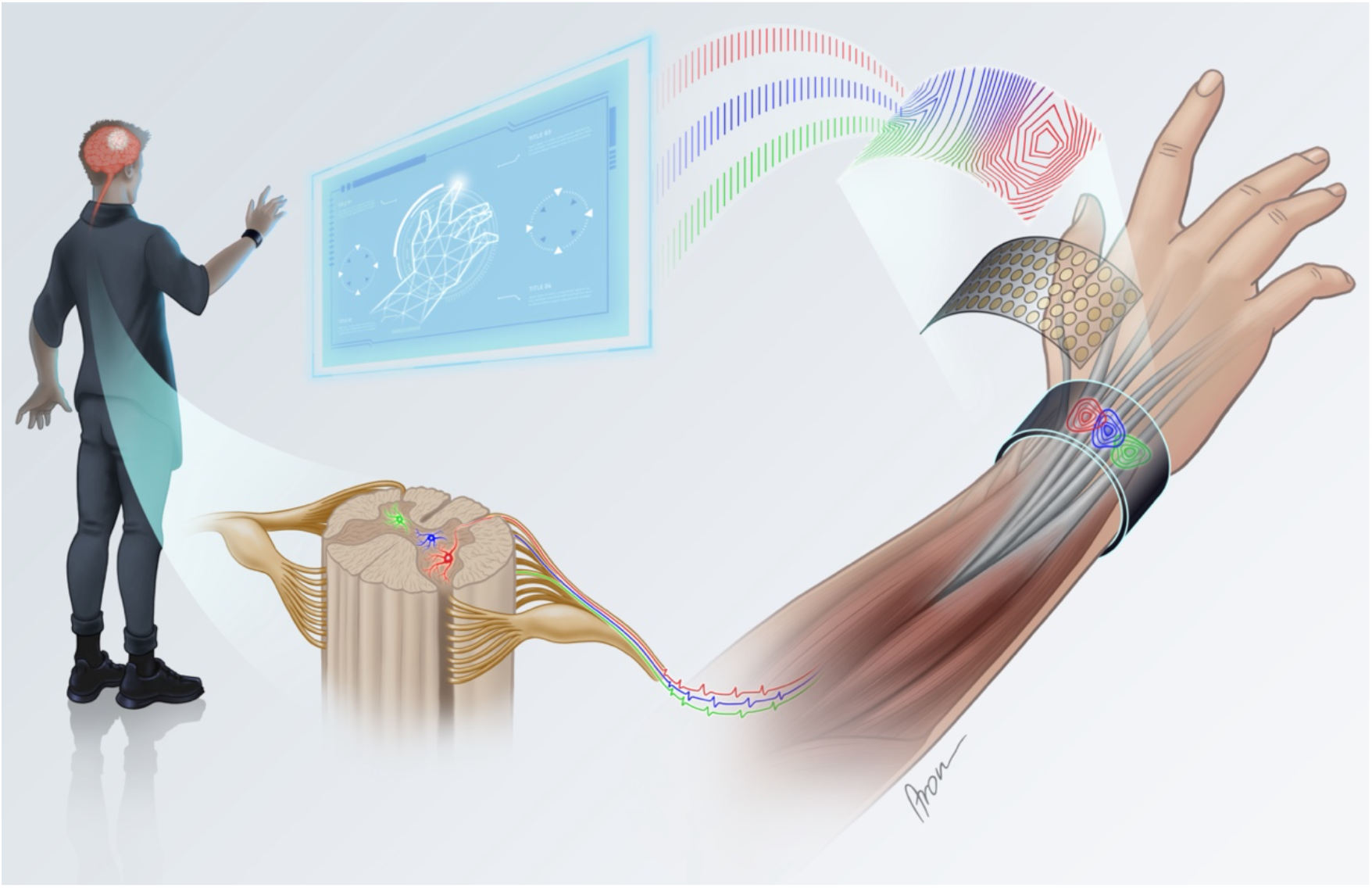
Interfacing motoneurons in the spinal cord non-invasively from the wrist via volume conduction and tendon potentials. Motoneurons in the spinal cord translate the synaptic inputs they receive from supraspinal centres and peripheral afferents into a neural output sent to the muscles. When excited, they discharge axonal action potentials that reach the neuromuscular junction of the innervated muscle fibers. The associated electric fields can be detected at the tendon endings of the wrist using electrodes (in this case, arranged in a wrist-band) due to volume conduction. However, the obtained tendon electric signals experience a high crosstalk between each other due to the convergence of multiple tendons in the reduced space of the wrist. To enhance the information transfer needed for precise decoding of motor volition, the generative model of the recorded potentials can theoretically be reversed to estimate the activity of the spinal motoneurons as long as the induced electric fields at the tendons are unique for each motoneuron.

A few studies have recorded the electrical activity at the wrist to decode motor intention(*14*–*16*), but the reported accuracy has been generally low. Using temporal EMG features, Botros et al.(*16*) achieved 88% classification accuracy in offline prediction of individual finger tasks and Jiang et al.(*15*) obtained 75% in a real-time control task. These accuracies are lower than those generally reported with classic EMG recordings from the forearm(*17, 18*). This poor performance has been explained by the convergence of multiple muscle tendons in a reduced space, which results in high crosstalk between the signals recorded at the wrist.

Estimating activity of spinal motoneurons that indirectly generate the recorded fields is an alternative approach for recording electric potentials at the wrist achieved by reversing the generative model of the recorded potentials (Fig. 1). This approach has been developed for muscle recordings but not for tendon potentials. For muscle signals, the inverse problem is often solved by blind source separation(*19*–*23*). The sparseness of the estimated sources is maximized under the assumption that the action potentials of the muscle fibers innervated by each motoneuron are unique with respect to those elicited by other motoneurons. The latter assumption is satisfied when the number of observations (sensors) is sufficiently large, i.e. in the range of tens to hundreds(*24*).

The electrical activity recorded over tendon tissue can also theoretically be separated into contributions of individual neural sources. Each motoneuron discharge generates an electric field transmitted through the tendon tissue that can be distinguished from the fields generated by other motoneurons. The biophysical properties of electric potentials recorded over tendon tissues are known (*25*–*28*), but these signals have not previously been used to identify individual motoneuron discharges.

Here we propose an innovative interface that decodes spinal motoneuron discharges from tendon electric signals at the wrist to develop a non-invasive, unobtrusive, and socially acceptable wearable. The scientific rationale of the approach is that the end-of-fiber components of the muscle fiber action potentials that produce the tendon electric fields have the same timing (with a constant delay of a few ms) as the axonal action potentials from the spinal cord(*29*). Therefore, we recorded tendon electric signals with modular and flexible electrode arrays that adapt to the users’ body(*30*). This compact design can also be paired with highly embedded systems and wireless communication to develop wearable recording devices(*31*). Here we demonstrate that the decoded tendon signals from these recordings indeed correspond to individual spinal motoneuron discharges, and that this neural decoding enhances the information transfer to accurately predict individual and combined finger movements in offline and real-time control conditions. With this demonstration, we present a new high-fidelity, non-invasive, unobtrusive, and socially acceptable wearable neural interface at the wrist as a viable alternative to invasive neural recordings or traditional muscle interfaces.

## RESULTS

To validate the wrist-wearable neural interface, we investigated the physiological properties of the decoded tendon electrical signals, and then assessed their potential for offline and real-time prediction of finger tasks. In experimental sessions, nine participants performed isometric contractions of individual fingers as well as of the combinations of thumb, index, and middle fingers at 15% and 30% of maximal force. During the tasks, high-density electrode arrays were placed around the circumference of the wrist (at least 100 channels arranged in rows of 5 electrodes; Fig. 2) to record tendon electric signals. Supplementary Fig. 1 shows the spatial distribution of the average activity recorded at the tendon for each finger flexion and subject. In addition, high-density electrode arrays were mounted along the circumference of the forearm (not shown in Fig. 2, see Supplementary Fig. 2) to validate the neural nature of the decoded tendon electrical activity (see Methods).

**Fig. 2.**
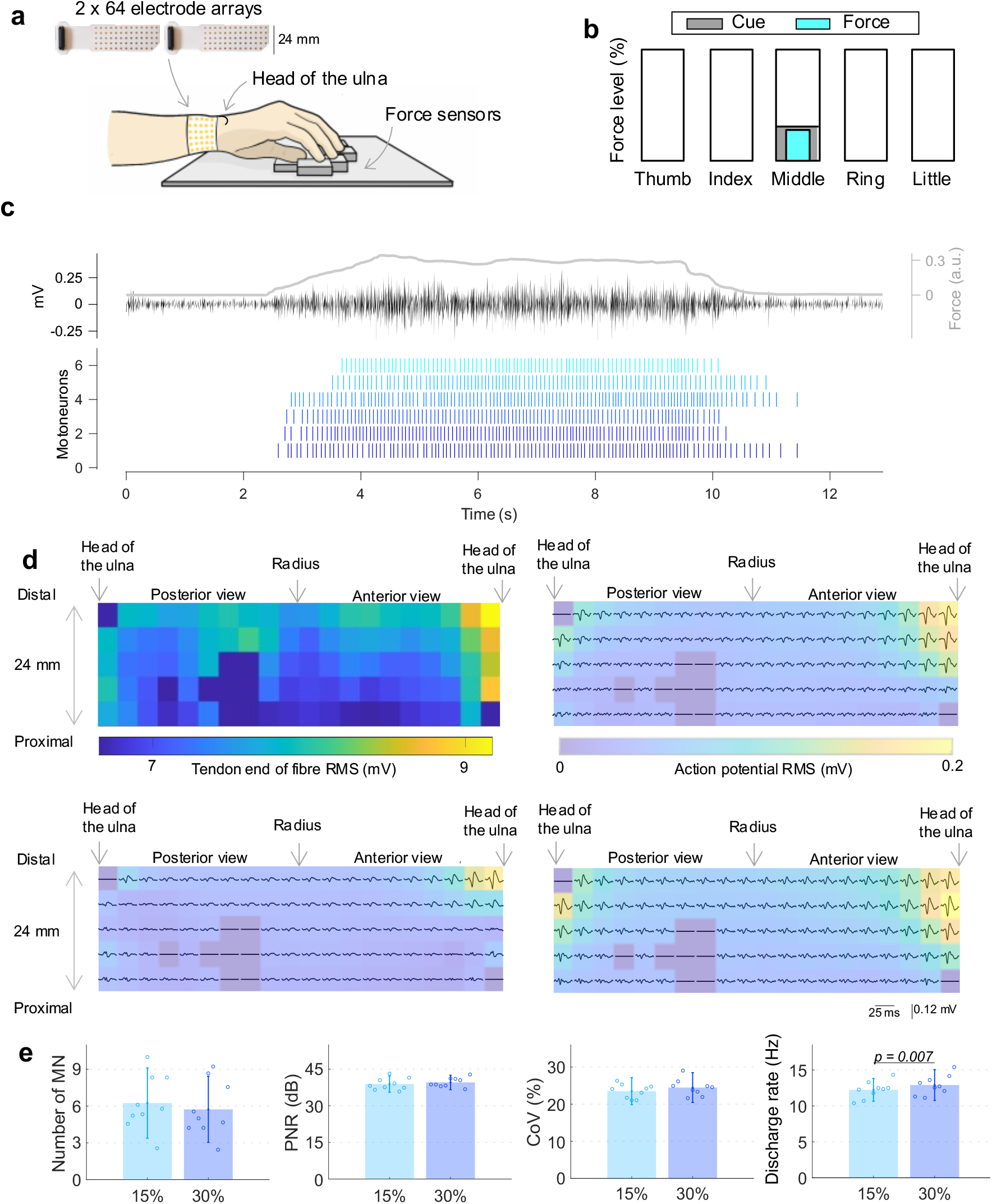
Decoded tendon electric signals from the wrist interface. (**A**) Experimental setup for the concurrent acquisition of the tendon electric signals and individual finger flexion forces. (**B**) Participant’s visual feedback with the contraction cues in grey and the exerted forces in blue. (**C**) A representative contraction from one participant with the force profile in grey, one tendon electrical signal from the tendons in black, and the decomposed spike trains (decoded tendon electrical signals). (**D**) (top left) 2D spatial distribution of the root mean square (RMS) of each channel of the tendon electrical signals from one representative contraction, (top right and bottom) three examples of the reconstructed action potentials of three decoded motoneurons from the same contraction after spike-triggered averaging along with their corresponding RMS spatial maps. The channels in dark blue were discarded due to noise interference. (**E**) Physiological analysis of the decoded tendon electrical signals from the wrist for 15% (light blue) and 30% (dark blue) force efforts in terms of the number of detected motoneurons (MN), pulse-to-noise ratio (PNR), coefficient of variation (CoV) of the inter spike intervals, and motoneuron discharge rate averaged across finger movements and subjects. The results indicate that the decoded tendon electrical signals from the wrist are accurate (PNR > 30 dB and CoV < 30%) and comply with motoneuron’s physiological behaviour (DR between 5-25Hz). The reported significance levels are based on two-way repeated measures ANOVA.

### Physiological analysis

The tendon electric signals were decomposed into a series of discharge timings (decoded tendon electric signals) by a convolutive blind source separation algorithm (see Methods). A representative contraction with the force profile, one tendon electric signal, and the corresponding decoded activity are depicted in Fig. 2.c. Once the tendon signals have been decoded, the spatial representation of the electric potentials can be recovered by spike-triggered averaging the tendon electric signals over time intervals centered at the detected discharge times (see Methods). Fig. 2.d shows representative 2D amplitude maps of tendon electric signals and three examples of the electric potentials generated by single motoneurons and volume conduction. The distributions of electric potentials recovered from the wrist are unique for each motoneuron, with high synchronization in their peak amplitude times due to the end-of-fiber nature of these electrical activities.

On average, 6 ± 3 motoneurons were identified by source separation of the tendon electric fields per finger contraction at each force level. To ensure that this decoded tendon electrical activity indeed represented the neural output from the spinal cord, the discharge timings of the decoded tendon electric signals were used to trigger an average of the EMG signals concurrently recorded at the forearm (see Methods and Supplementary Fig. 2). The rationale for this processing is that if the discharge times decoded at the wrist correspond to the times of activation of spinal motoneurons, then the triggered average should identify muscle fiber potentials at the forearm above the baseline noise. Indeed, motoneuron activity determines muscle fiber activity synchronous with the motoneuron firings. Therefore, if the decoded times of activation from the wrist determine action potentials of muscle fibers at the forearm when used as triggers, they must correspond to discharge patterns of motoneurons. This approach provided a means for robustly validating the wrist neural interface. The action potentials at the forearm obtained by spike-triggered average were considered above the baseline noise if their peak amplitude was greater than four times the noise level, as commonly assumed in spike sorting(*32*) (see Methods and Supplementary Fig. 2). From the total population of 970 detected motoneurons at the wrist, 703 (72.47%) resulted in detectable action potentials at the forearm. This is an extremely high proportion, considering that motoneurons detected at the wrist may innervate muscle fibers deep into the muscle which therefore would not produce sufficiently large action potentials at the forearm skin surface. This result indicated that the timings of activation decoded from the tendon electric fields indeed correspond to neural activity from the output layer of the spinal cord. This demonstrates that a peripheral recording from the skin overlying tendon tissue can be decoded into the ultimate neural code of movement.

The quality of the decoding of tendon electric fields was further validated with a measure of pulse-to-noise ratio(*33*) (PNR) of the estimated discharge activation patterns. The PNR is an estimate of the mean square error of the motoneuron spike detection that measures the ratio between the mean energy of the spikes at the discharge times and the baseline of the signal(*33*). At both force levels, the PNR was greater than 30 dB and generally higher than usually observed when decoding classic EMG recordings from the forearm(*34*–*37*) (38.9 ± 2.3 dB and 39.6 ± 3.0 dB for 15% and 30% force efforts, respectively) (Fig. 2.e). These levels of PNR correspond to an accuracy in detection of spikes in the estimated sources with >90% sensitivity and < 2% false alarm rate(*33*).

After validating the decoding procedures, we further analyzed the properties of the decoded discharge patterns to verify whether they were consistent with known physiological properties. We extracted the average discharge rate of each identified motoneuron as well as the coefficient of variation of the estimated inter-spike intervals (ratio between the standard deviation and mean of the inter spike intervals expressed as a percentage). The estimated motoneuron discharge rates were within the physiological range of 5-25 Hz(*38, 39*) at both force levels (12.23 ± 1.58 Hz and 12.90 ± 2.14 Hz at 15% and 30% force efforts, respectively) (Fig. 2.e), being significantly higher at 30% than at 15% force effort (F_1, 8_ = 12.879, p = 0.007), in agreement with motoneuron’s rate coding in force production(*9*). The coefficient of variation of the estimated inter spike intervals was 23.51 ± 3.63% and 24.47 ± 4.01%, for 15% and 30% force efforts, respectively (Fig. 2.e), which is within known physiological values(*40*).

The analysis of accuracy via spike-triggered average and PNR, as well as the physiological analysis of motoneuron behavior demonstrated the validity and accuracy of the proposed decoding technique. Overall, these results prove the accurate identification of the activity of individual spinal motoneurons through non-invasive wearable recordings overlying the tendon endings at the wrist. After confirming validity and accuracy, we established a human-machine interface based on the proposed neural decoding approach.

### User intent prediction (offline)

The decoded motoneuron activity was used to classify finger movements (Fig. 3). As a reference, we compared the classification from motoneurons with that obtained using the tendon electric signals before decoding. Relevant features for pattern recognition (see Methods) were extracted from the motoneurons and tendon electric signals and were fed into independent neural networks for classification. In a first scenario, the neural network was trained with the steady contraction part of all finger tasks and rest (10 classes in total) at either 15% or 30% force effort following a ten-fold cross validation approach (see Methods and Fig. 3). Figure 3 shows the resulting classification accuracy, which was significantly higher for the decoded motoneurons than for the un-decoded tendon electric signals at both 15% (96.93 ± 2.09 % vs 81.23 ± 10.04 %; F_1,8_ = 23.379, p = 0.001) and 30% force efforts (97.60 ± 1.75% vs 85.62 ± 6.86 %; F_1, 8_ = 31.036, p = 0.001). In addition, the effect of force was only significant for the tendon electric signals, yielding in higher classification accuracy at the highest force effort (F_1, 8_ = 8.026, p = 0.022).

**Fig. 3.**
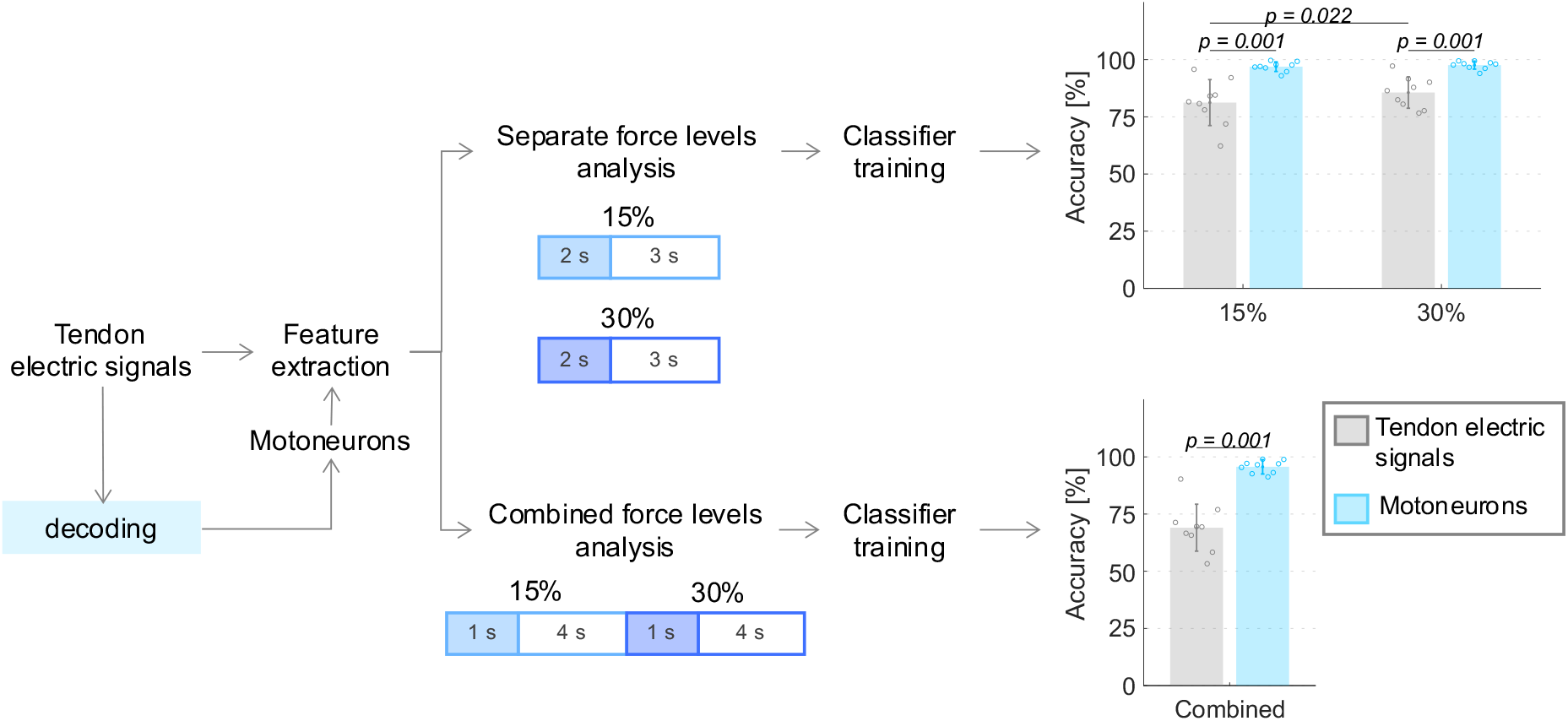
Offline classification performance. Processing steps for the tendon electrical signals and decoded motoneurons from the wrist for the separate and combined force level analyses for finger prediction. The blocks represent the steady contraction part of the signals with the training and testing portions in blue and white, respectively. The obtained classification accuracies show that the decoded motoneuron activity (in blue) is a better predictor of underlying finger flexion than the un-decoded tendon electric signals (in grey), with high accuracy, irrespective of the force level. Plot significances are based on repeated measures ANOVA.

To simulate more realistic control conditions with variable force levels, the classification accuracy was also calculated after training and testing with finger contractions from both force levels combined following a ten-fold cross validation (see Methods and Fig. 3). The gain in classification accuracy when decoding the tendon signals was even greater in this condition (95.65 ± 2.76 % vs 69.04 ± 10.61% for decoded and un-decoded tendon signals, F_1,8_= 64.606, p < 0.001). The results obtained without decoding the neural activity were consistent with those reported in previous studies(*15*) and indicate poor classification performance. Conversely, the proposed neural decoding allowed for >95% accuracy over ten finger tasks at multiple force levels, which was substantially greater than without decoding the tendon signals as well as than conventional EMG-based interfaces(*16, 17, 41*).

Overall, the motoneuron activation patterns identified from the wrist provided a highly accurate prediction of finger tasks.

### Real-time control

We then implemented the decoding and classification in real-time and tested the resultant interface in an online control task on four participants. As for the offline analysis, we compared the real-time control results with the control achieved without decoding the tendon electric signals. Figure 4a shows the processing pipeline. Three repetitions of rest, plus each individual finger contraction, and all combinations of thumb, index, and middle (10 tasks in total) were recorded to train a neural network using the same signal features as for the offline analysis (see Methods and Supplementary Fig. 3). The training set was also used to calibrate the decoding parameters to be thereafter applied in real time to extract the corresponding motoneuron activity (see Methods). The motoneuron decoding and the classification were then applied online and used for control of finger tasks. Figure 4b shows the decoded signals obtained during this process from one representative participant. On average, 78 ± 8 motoneurons were identified across all tasks for each participant. During the online tests, 4 targets per class (40 in total) were presented to the participants in randomized order. The participants were given 5 s to attempt each target with a success condition of maintaining the correct gesture for 500 ms (Fig. 4c). The mean completion rate resulted significantly higher for the motoneuron (93.12 ± 2.39 %) than for the un-decoded tendon electric signals (56.87 ± 18.41%), while maintaining similar completion times (1.81 ± 0.89 s vs 1.65 ± 0.82 s).

**Fig. 4.**
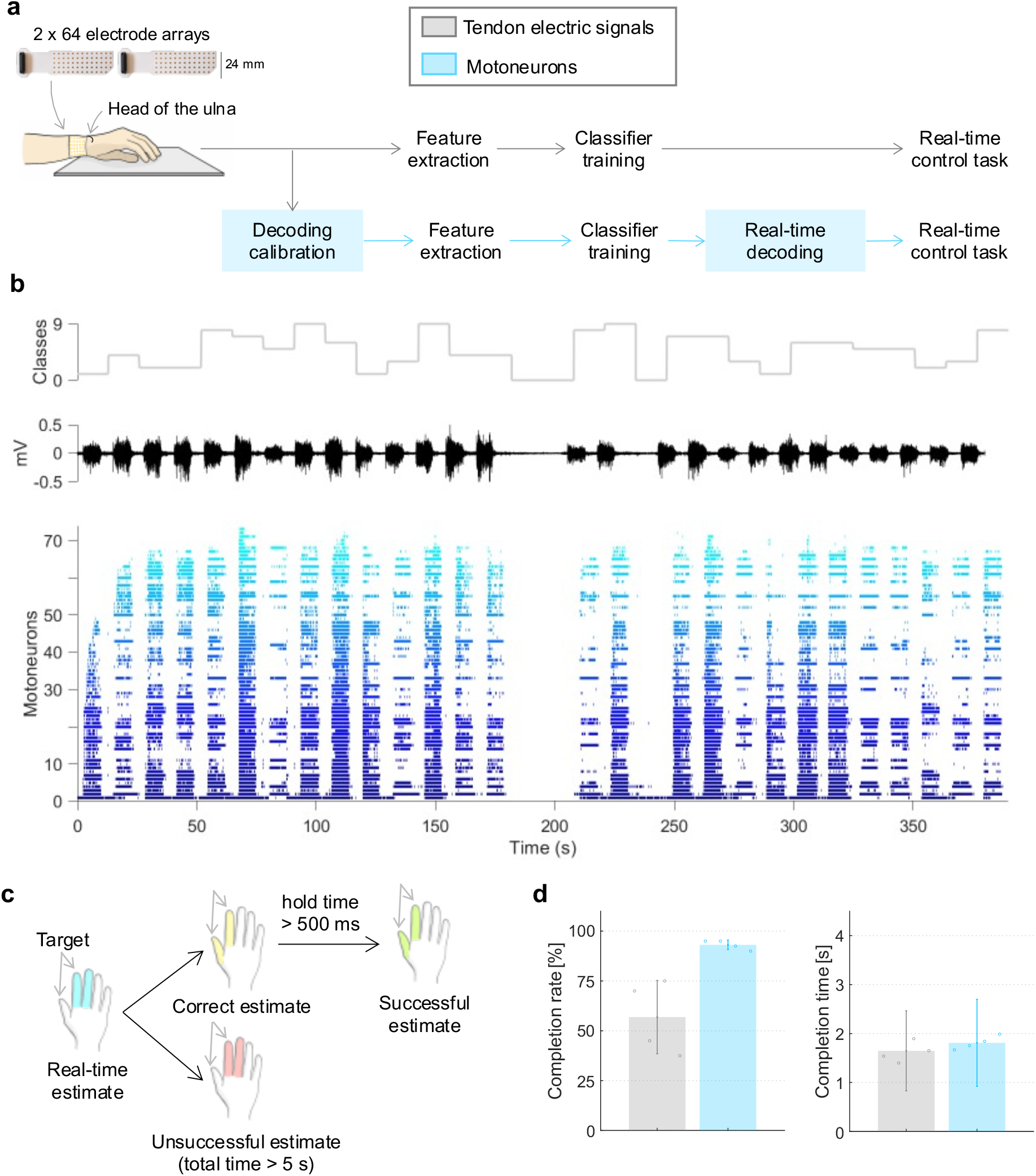
Real-time control. (**A**) Acquisition setup for the online testing with the processing pipelines for the tendon electric signals (grey) and decoded motoneurons (blue). The additional steps specific of the decomposition algorithm are highlighted in blue. (**B**) Training set from one representative participant with the class cues on top (each individual finger plus all the combinations of thumb, index, and middle), one tendon electric signal in the middle, and the decoded motoneuron activity during the decomposition calibration at the bottom. (**C**) Success and fail conditions for the real-time control task (**D**) Online control performance for the tendon electric signals (grey) and decoded motoneurons (blue) in terms of completion rate and completion time.

The poor online control capacity when using tendon electric signals from the wrist without decoding is in agreement with previous work(*15*) and indicates the poor discrimination power of mixtures of electric fields activated by motoneurons. Conversely, the control using separated motoneuron activation patterns was extremely accurate and provided large information transfer (10 classes, ∼93% successful task completions).

## DISCUSSION

We have demonstrated that the neural information sent from the spinal cord to muscles can be accurately decoded at the single motoneuron level with a wearable technology mounted at the wrist. This technology allowed for the accurate real-time control of 10 commands elicited by finger tasks. Thus, we have designed a unique neural interface with high information transfer rate. The presented results show the potential of future portable, battery-operated systems worn at the wrist as un-obtrusive and viable neural interfaces to use in daily living.

The neural origin of the decoded tendon signals was proven by retracing the muscle fiber action potentials from concurrent recordings from the forearm via spike-triggered averaging (Supplementary Fig. 2). This approach showed that the majority of the sources identified at the wrist coincided with action potentials at the forearm muscle fibers which were well above the baseline noise. This result demonstrates the neural origin of the decoded activity. Indeed, if the decoded activity were not generated by motoneurons, spike-triggered averaging on muscle electrical signals would yield only noise as it would be equivalent to average uncorrelated EMG signals at the forearm. The observation that approximately 30% of the decoded motoneurons did not result in averaged potentials above the noise level is explained by the location of the muscle fibers innervated by the detected motoneurons. For instance, the flexor digitorum superficialis, the flexor digitorum profundus, and the flexor pollicis longus are located deep in the forearm and therefore their innervating motoneurons generate electric potentials detectable at the wrist that may not be at the forearm. Interestingly, this indicates that the limitations of EMG recordings to superficial muscles may be surpassed by tendon recordings when multiple muscles converge into a common tendon area, such as at the wrist. In addition to proving the neural origin of the decoded activity by spike-triggered averaging, we also computed the pulse-to-noise ratio (known estimate of the mean square error(*33*)), coefficient of variation of the inter spike intervals (associated to the likelihood of erroneously detected action potentials(*40*)), and action potential discharge rate (used as a physiological indicator(*38, 39*)). All metrics were well within the expected accuracy and physiological standards, showing that the decoded tendon electric signals were reliably extracted and corresponded to the neural output from the spinal cord (Fig. 1).

The number of identified motoneurons (6 motoneurons per finger contraction at both force efforts) was consistent with the results by Stachaczyk et al.(*42*) who identified between 5-8 motoneurons per finger contraction when recording signals from the forearm flexor muscles. Interestingly, however, the pulse-to-noise ratio levels observed at the wrist in this study were greater than those usually reported for forearm recordings. Moreover, as discussed above, the decoding from the wrist was not biased towards detecting superficial muscles since the effect of the volume conductor at the wrist is less than at the forearm. Furthermore, the signal characteristics at the wrist are different than that at the belly of the muscle. The electric potentials recorded at the wrist are end-of-fiber components(*25*), which are non-propagating potentials highly synchronized across channels (as shown in Fig. 2d) and shorter in time than the propagating signals recorded from muscles(*43*). These characteristics result in greater temporal sparsity at the tendons because of the shorter duration of the individual potentials. Therefore, the raw tendon signals can be modelled as convolutive mixtures in which the filters applied to the sources have relatively short duration (see Methods). Short duration filters are easier to compensate since they better approximate delta functions that represent the sources. This also results in sparser observations. Overall, we have not only shown that electrical potentials recorded from tendon tissue can be decoded into neural activity but also that recording from the wrist may even be preferable over conventional muscle recordings in term of representativeness of the decoded information and accuracy.

From the decoded signals, we performed an offline classification with the aim of using the wrist interface for control applications. Results showed that the decoded signals accurately predicted finger tasks for up to 10 classes with significantly higher accuracy than the tendon electric signals without decoding. The classification accuracy for the un-decoded tendon electric signals (∼83%) was slightly higher than the one obtained by Jiang et al.(*15*) in a real-time gesture prediction task (∼75%). Although only four sensors were used in that previous study, their location was consistent with our electrode placement below the head of the ulna. In contrast, Botros et al.(*16*) reported higher accuracies (∼88%) for offline single and combined finger prediction using the same feature set, but their electrodes targeted the muscle fibers in the proximal and medial part of the wrist instead of the tendons. The main limitation of the tendon electric signals is their high crosstalk due to the convergence of the muscle tendons in a reduced space. This contributes to the high overlap between the classes in the spatial activity maps (Supplementary Fig. 1) and feature space (Supplementary Fig. 3) of the tendon electrical signals, which resulted in an overall lower performance than the decoded signals. In contrast, previous studies by Dai and Hu showed that myoelectric activity spatial maps from the forearm can indeed differentiate between individual finger flexions(*44*) and extensions(*45*) when a large muscle area is covered. For the flexors, the performance even improved when both myoelectric and neural spatial maps were used(*44*).

To increase the information transfer and enhance the separability between finger classes despite the limited area, the tendon electrical signals need to be decoded into the neural output of the spinal cord. However, no other study has previously addressed the potential of the wrist for non-invasive neural interfacing, thus the only comparative results are from the forearm. In a finger prediction task, Stachaczyk et al.(*42*) obtained similar classification accuracy for the neural output of the spinal cord at the forearm (98%) to the presented here at the wrist (∼97%), although only for individual finger tasks (the four digits, excluding the thumb, while here we tested classification over 10 tasks comprising individual and combined finger gestures). Therefore, the reported accuracy in finger task classification when decoding motoneurons from the wrist is even superior to that of motoneurons decoded from muscle tissue. Stachaczyk et al.(*42*) also found that the neural output was robust to variations in the force level, unlike myoelectric signals from the forearm when predicting finger flexions(*42*). Indeed, the increased classification error of tendon electric signals when both force levels were combined was in agreement with previous literature on the effect of dynamic contractions in myoelectric pattern recognition(*46, 47*). These findings suggest that motoneurons discriminative power between fingers does not rely on spatial information, nor on force encoding. This is supported by the feature map of the decoded tendon electric signals presented in Supplementary Fig. 3 where the different classes exhibit higher separability than in the raw feature space, despite targeting the same area and corresponding to multiple force levels.

Additional real-time experiments were carried out with multiple repetitions to validate and extend the offline results to real-life interfacing scenarios. This analysis showed that the decoded tendon electric signals from the wrist can be accurately detected, and enabled real time interfacing with over 70 motoneurons. Moreover, this neural decoding led to high reproducibility and separability between finger contractions, as evidenced by the high task completion rate (>93%) in relatively short time (∼1.81s per task). To the best of our knowledge, no other study has previously implemented a pattern recognition approach with neural decoding in real time. In conclusion, we have shown the feasibility of accurate, non-invasive, real-time, neural interfacing with wearable sensors mounted at the wrist. These innovative results open an important perspective in neural interfacing for large scale applications, in medical devices and consumer electronics applications.

## MATERIALS AND METHODS

### Offline experiment

#### Experimental setup

Nine healthy participants (4 females, 5 males, ages: 23-31) volunteered in the study. Both informed consent forms and experimental protocols were approved by Imperial College London ethics committee in accordance with the Declaration of Helsinki.

Two flexible EMG electrode grids (64 channels arranged in 5×13 with 8 mm distance, GR08MM1305, OT Bioelettronica) were placed along the circumference of the wrist right below the head of the ulna by visual inspection and physical palpation. In addition, myoelectric signals were concurrently recorded from the circumference of the thickest part of the forearm using three EMG electrode grids (64 channels arranged in 8×8 with 10 mm distance, GR10MM0808, OT Bioelettronica). Both signals were simultaneously acquired by a multi-channel amplifier (Quattrocento, OT Bioelettronica), bandpass filtered between 10-500 Hz, and sampled at 2048 Hz with 16-bit ADC precision. Individual finger flexion forces were recorded concurrently at 10 Hz with 5 micro load cells (0-5kg CZL635, Phidget), located in an ergonomic and adjustable platform. The latter was designed to keep the hand supported while in a relaxed position. A custom program (Matlab 2019b, The MathWorks, Inc) was implemented to synchronously acquire all the signals.

Participants were seated on a chair with their arm supported and the fingers placed in a comfortable position on top of each force sensor. They were facing a computer screen where cues and visual feedback of their fingers’ flexion forces were provided. The maximum voluntary force effort across each finger was calibrated for each participant at the beginning of the experiment. Thereafter, they were instructed to follow the displayed trapezoidal cues (2 s rest, 2 s ramp up, 5 s steady contraction, 2 s ramp down and 2 s rest) at 15% and 30% of the maximum force effort for each individual finger and the combinations of thumb-index, thumb-middle, index-middle and thumb-index-middle in a randomized order (18 trials in total).

After the acquisition, tendon and myoelectric signals were digitally band-pass filtered between 20-500 Hz (zero-phase 20^th^ order Butterworth) and noisy channels (mostly by electrode overlap due to the excessive length of the electrode matrices) were discarded. On average 102 ± 12 channels were used for further analysis.

#### Decoding algorithm

Each axonal action potential of a motoneuron determines the generation of action potentials in the innervated muscle fibers. Once excited, the muscle fiber undergo depolarization in a confined portion of their membrane. The depolarization zone, which has a length of 5-10 mm, propagates along the muscle fibers from the end-plate to the tendons. At the tendons, the depolarization zone extinguishes, generating a so-called end-of-fiber potential. The electric signals recorded over tendon regions, therefore, are dominated by the end-of-fiber potentials. Interestingly, each end-of-fiber potential corresponds to an axonal action potential, such that the activation of motoneurons is reflected by electric fields generated at the tendons. For tendon regions that correspond to the convergence of tendon endings of multiple muscles, such as at the wrist, the recorded signals are the combination of end-of-fiber potentials from multiple muscles. These tendon regions therefore are bio-screens for the multiple pools of motoneurons innervating several muscles (Fig. 1), whose coordination generates movements. Interestingly, the end-of-fiber potentials have temporal high-frequency components and are less attenuated by the volume conductor than muscle propagating potentials(*48*). These properties make recordings at the tendons not only a feasible but also a more suited solution for neural decoding than direct muscle recordings (see Discussion).

Given the above description of the origin of electric potentials recorded at the tendon regions, the following mathematical model holds:

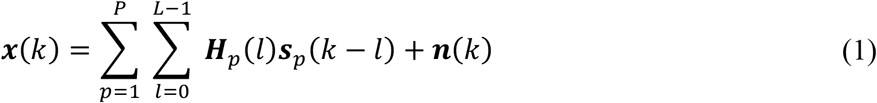

where ***x***(*k*) are the tendon electric signals at time *k* generated by the additive contributions of *p* spinal motoneuron pools innervating different muscles, plus independent noise ***n***(*k*). The activity of individual motoneurons is modelled as trains of delta functions at their corresponding discharge timings, convolved by their respective end-of-fiber potentials along their duration *L*. In equation (1), ***s***_*p*_ and ***H***_*p*_ represent the delta trains and end-of-fiber potentials of all the motorneurons in the *p*^th^ pool, respectively. The high-frequency nature of the end-of-fiber components at the tendons corresponds to short temporal durations compared to the propagating ones, which translate in relatively low values for *L*. In this way, the end-of-fiber potentials (***H***) better approximates to the source delta functions than muscle fiber potentials. In addition, equation (1) also shows that the high temporal sparsity of the end-of-fiber potentials is also reflected in the mixed tendon electric signals (***x***(*k*)).

Although the previous formulation provides a clear interpretation of the relation between the neural and volume conductor elements of the model, it complicates the de-mixing of its components. To simplify it, the matrices of the end-of-fiber potentials (***H***_*p*_) and motoneuron firings (***s***_*p*_) can be rewritten including their corresponding delayed versions along *L* to compensate for the effect of the convolution. Moreover, tendon electric signals (***x***(*k*)) should also be extended to an artificial delay proportional to *L* and inversely proportional to the number of electrodes, to offset the increase of motoneurons to estimate. This yields the following equation:

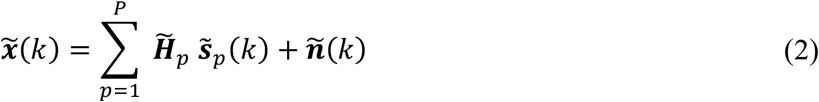

where ∼ indicates the extended variables. Equation (2) reflects the presence of multiple spinal motoneuron pools innervating different muscles due to the convergence of their tendon endings at the wrist. Nevertheless, their estimation can be conveniently carried out in a single matrix form by concatenating the contributions of each pool to the global end-of-fiber potentials 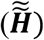 and motoneuron firings 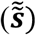 as follows:

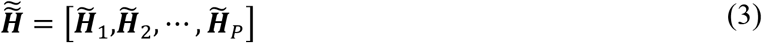

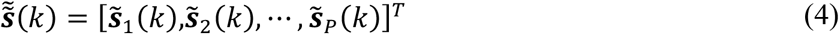

Such that:

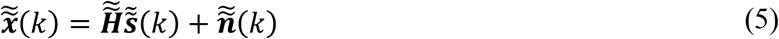

This expression can then be inverted by applying linear-instantaneous blind source separation while maximizing the sparseness of the sources, as long as the tendon potential generated by each motoneuron is unique in relation to the potentials generated by other motoneurons. This condition has been extensively validated for the propagating components of the muscle fiber action potentials(*37, 42, 45, 49*), but it has never been tested for the end-of-fiber components generated at the tendon endings. Therefore, this assumption needed to be confirmed experimentally (see Results).

To validate it, the convolutive blind source separation(*22*) algorithm was used to invert the end-of-fiber potentials of the model (finite impulse response filters) and decode the motoneuron firings from the recorded tendon electric signals. Briefly, convolutive blind source separation applies an initial whitening and fast fixed-point algorithm that maximizes sparseness(*50, 51*) to detect the unique sources (motoneurons), followed by a peak-detection and clustering postprocessing to identify their corresponding discharge timings in the estimated delta trains(*22*). After this process, only the original motoneuron firings (non-delayed versions) were kept for the rest of the analyses. Finally, for the offline analysis only, the output of this fully automatic decomposition was validated in a semi-supervised approach that enables the modification of the thresholds of the local peak detection algorithm to update the filters of poorly detected sources and recalculate the motoneurons firings(*52*). Repeatedly detected motoneurons within each contraction (with > 30% shared spike timings(*40*)) were removed at this stage.

The decomposition output was evaluated in terms of the number of identified motoneurons at the wrist and the percentage of those that corresponded to electric potentials occurring concurrently at the forearm. To do so, the fiber potentials at the forearm were calculated by spike-triggered averaging the forearm signals in 50-ms windows using the discharge times identified from the wrist as triggers. The average potentials obtained in this way were considered detectable if their peak amplitude was higher than four times the standard deviation of the baseline noise(*32*) computed over the first and last 15ms of the spike triggered average. If one or more channels in the array met this condition, it was concluded that the corresponding source identified from the wrist corresponded to the activation of muscle fiber action potentials and thus to the discharges of a spinal motoneuron.

The accuracy of the decoding was assessed based on the pulse-to-noise ratio of the estimated spike train (mean of the detected spiking activity divided by the mean baseline of the estimated source expressed in dB(*33*)). In addition, the coefficient of variation of the inter spike intervals (ratio between the standard deviation and mean of the inter spike intervals expressed as a percentage) and motoneurons’ discharge rate (ratio between the number of action potentials fired by a motoneuron and their active period measured in seconds) were computed to evaluate their physiological properties.

A two-way repeated measures ANOVA was used to evaluate differences in the number of motoneurons, pulse-to-noise ratio, discharge rate, coefficient of variation between force levels and finger flexions. Statistical significance was set to p < 0.05 and all calculations were performed in IBM SPSS Statistics 26. Normal distribution of all variables (9 fingers x 2 force levels) was verified by the Shapiro-Wilk test of normality (p > 0.05). Few exceptions were found in 1) the number of motoneurons in the little finger at 15% force effort (p = 0.019), 2) the pulse-to-noise ratio of the thumb at 15% (p = 0.037) and index at 30% (p = 0.005), and 3) coefficient of variation of the little at 15% (p = 0.006), thumb-index at 15% (p = 0.029) and thumb at 30% (p = 0.001). However, this low proportion of non-normal levels was considered acceptable for the two-way repeated measures ANOVA. The assumption of sphericity was checked for the finger flexion factor (levels > 2) by Mauchley’s test and if not satisfied, the Greenhouse-Geisser correction was applied to the degrees of freedom. Since no two-way interaction between the factors was found (p > 0.05), the main effects of force level and finger flexion were analyzed by one-way repeated measures ANOVAs. Bonferroni correction was applied for pair-wise comparisons between finger flexion levels.

#### Task classification analysis

For the control analysis, motoneurons across contractions were tracked based on the 2D correlation coefficient between their motoneuron action potential maps. These maps were calculated by spike-triggered averaging over 25-ms windows of all raw tendon signals at the wrist electrode array centered at the timings of the motoneurons’ spikes(*53*). The analysis was carried out only for those channels with significant peak amplitude (i.e. higher than two times the standard deviation of that motoneuron’s peak amplitudes among all channels). Motoneurons were considered the same if their normalized cross-correlation coefficient exceeded 0.70.

Thereafter, the steady contraction part (5s) of each finger flexion was selected and concatenated along with 5s of rest for feature extraction. Motoneurons firings were windowed in intervals of 120 ms with 40-ms step to compute the spike count of each motoneuron. Raw tendon electric signals were windowed alike to extract four time-domain features(*54*) (root mean square, slope sign changes, zero crossings and waveform length) for each channel. Although multiple features have been proposed to decode movement intentions from electric signals(*16*), the selected feature set is the most common in pattern recognition tasks(*55*).

A multilayer perceptron with one hidden layer and ten hidden neurons(*56*) was used to classify the raw and decoded tendon features separately into 10 classes (9 finger flexions plus rest). The multilayer perceptron was trained using the gradient descent with momentum and adaptive learning rate backpropagation algorithm. Performance was evaluated in terms of classification accuracy applying ten-fold cross-validation. In addition, classification accuracy was calculated for two scenarios: training and testing with contractions from one force level only, and from both levels combined. In both cases, 2 s of each class were used for training and the remaining data (3 s for the single force dataset, and 8 s for the combined) for testing.

In the separate case, a two-way repeated measures ANOVA was used to evaluate differences in classification accuracy between force levels and data type. Statistical significance was set to p < 0.05 and all calculations were performed in IBM SPSS Statistics 26. Normal distribution was validated by the Shapiro-Wilk test of normality with the only exception of decoded tendon signals accuracy for 15% force level (p = 0.004). When a statistically significant two-way interaction was found between force levels and data type (F_1,8_ = 13.807, p = 0.006), the simple main effects were analyzed with focused one-way repeated measures ANOVA fixing the levels of the interacting factors. When both force levels were combined during the training and testing, a one-way ANOVA was used to assess differences in classification accuracy due to the data type (2 levels). Normal distribution was again ensured by the Shapiro-Wilk test of normality.

### Real-time experiment

Four healthy participants (2 females, 2 males, ages: 25-32) participated in the online experiment (approved by Imperial College London ethics committee in accordance with the Declaration of Helsinki) after signing informed consent forms.

In this case, only tendon electric signals from the wrist were acquired following the same setup previously described. Participants were comfortably seated in front of a computer screen, with their right hand resting in a neutral position on top of a table. During training, they were asked to perform isometric finger contractions against the table at up to a comfortable level following trapezoidal cues (2 s rest, 2 s ramp up, 5 s steady contraction, 2 s ramp down and 2 s rest). Three repetitions each individual finger, the combinations of thumb-index, thumb-middle, index-middle and thumb-index-middle, as well as rest, were recorded in a randomized order (30 trials in total).

To extract motoneurons’ firings in real-time, we implemented a dual phase blind source separation(*23*). In the first calibrating phase, the algorithm followed the same procedure described above to identify the latent motoneurons. Then, the obtained inverse of the end-of-fiber potential filters was applied to new tendon electric signals epochs to decompose the activity of the previously identified motoneurons, and detect new action potentials using the stored spike and noise centroids of each source(*23*). During calibration, motoneurons with more than 20% of shared spikes were considered equal and only the one with highest pulse-to-noise ratio was preserved to avoid redundant activity. In this case the training set was used to calibrate the decomposition parameters for its later implementation in real-time during the control task. As in the previous experiment, the spike count of each motoneuron was calculated over sliding windows of 120 ms with 40 ms step (coinciding with the update rate of the system). On the other hand, the root mean square, slope sign changes, zero crossings and waveform length were extracted for the raw tendon electric signals using the same windowing process.

Two multilayer perceptron with one hidden layer and ten hidden neurons(*56*) were used to classify the decoded and raw tendon features separately into 10 classes (9 finger flexions plus rest). They were trained using gradient descent with momentum and adaptive learning rate backpropagation over the steady part of the contraction. Both multilayer perceptrons were tested separately in a real-time task with 4 targets of each class (40 targets in total). Participants were given 5 s to attempt each target, with a required hold time of 500 ms to consider the target successfully reached. Performance was measured in terms of completion rate (number of successful targets divided by the total number of targets, expressed as a percentage) and completion time (time needed to successfully achieve a target).

## Funding

Engineering and Physical Sciences Research Council, Centre for Doctoral Training in Neurotechnology for Life and Health (I.M.)

Facebook (I.M, D.Z.W)

Academy of Finland, project “Hi-Fi BiNDIng”, no. 333149 (I.V.)

European Research Council, Synergy grant, project Natural BionicS, no. 810346 (D.Y.B., D.F.)

## Author contributions

All authors conceived the study

I.M. performed the data acquisition.

I.M. conducted the analysis.

All authors interpreted the data.

All authors wrote and edited the manuscript

## Data and materials availability

All data are available in the main text or the supplementary materials.

## SUPPLEMENTARY MATERIALS

**Fig. S1.**
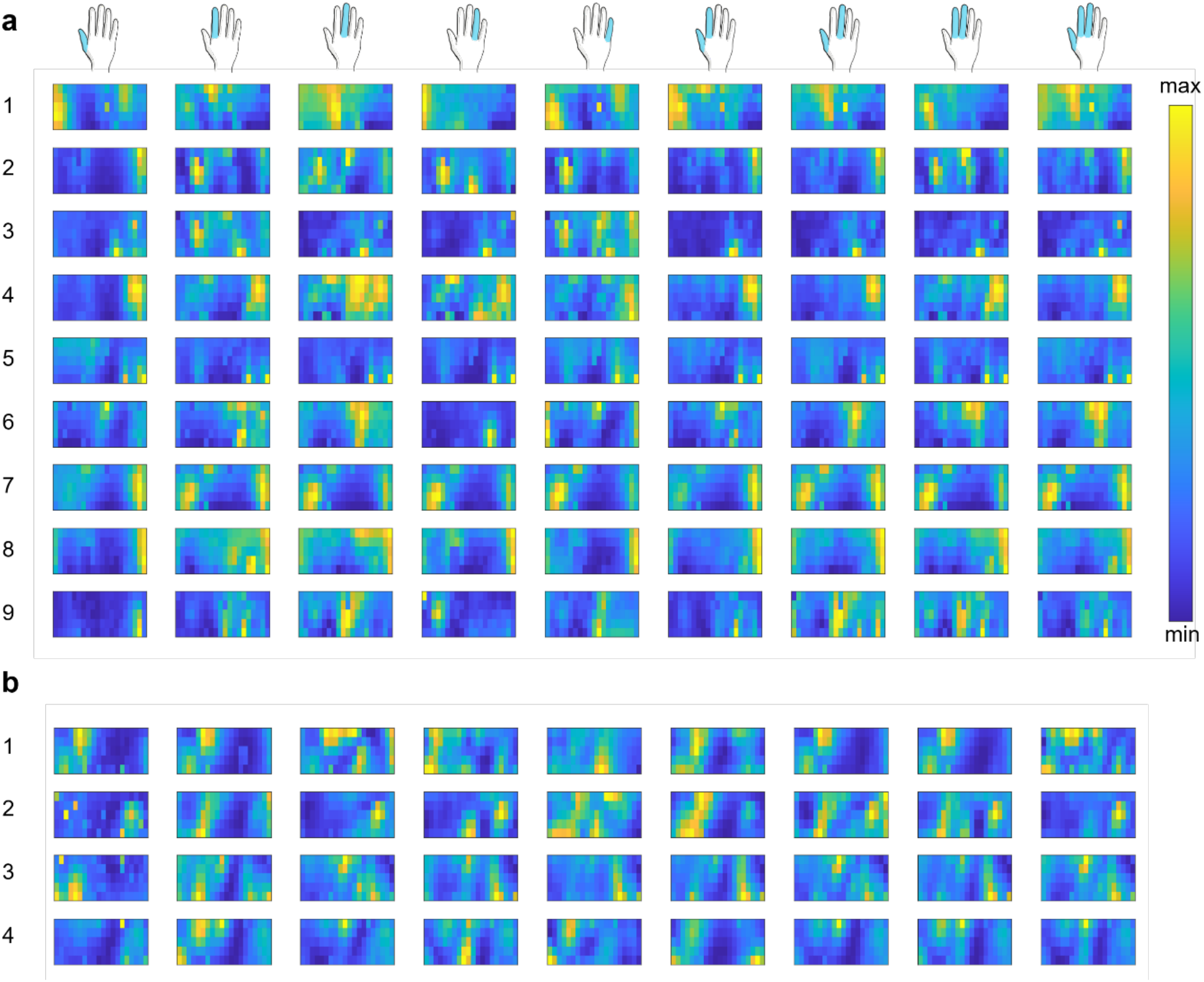
2D spatial distribution of the normalized amplitude of the tendon electric signals. Each subplot depicts the normalized root mean square of the tendon electric signals for each channel (pixel) in their corresponding spatial distribution in the electrode array at the wrist (view: bottom = proximal, top = distal, left = ulna posterior, right = ulna anterior). The values for few discarded channels due to noise have been estimated by 2D linear interpolation. The figure shows a high overlap in the activity area of the different finger contractions (columns) within each subject (rows). (**A**) mean across the maps at 15% and 30% of force efforts from the offline dataset. (**B**) mean across the maps of the three repetitions of the training set for the online prediction task.

**Fig. S2.**
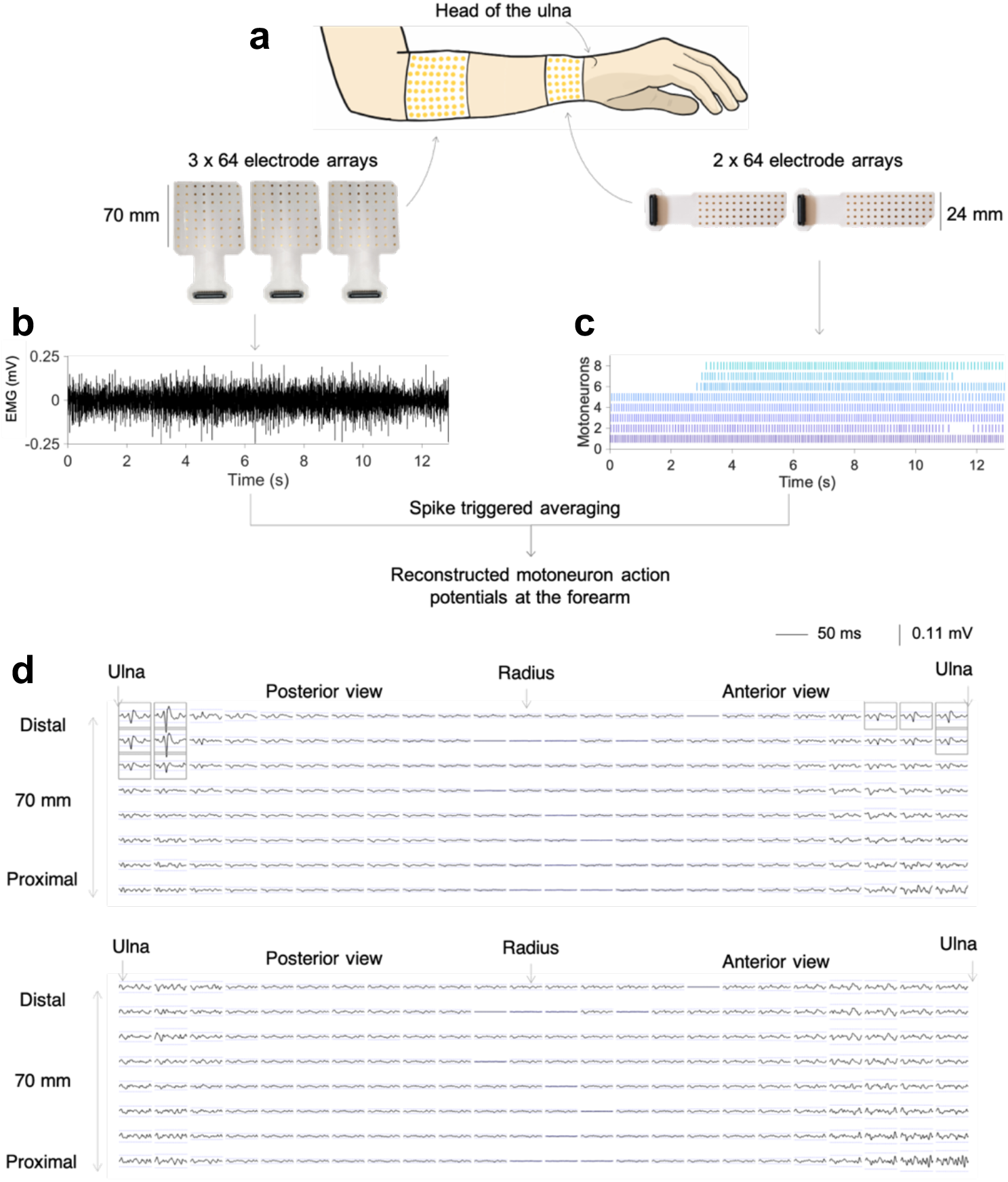
Retracing motoneuron fiber action potentials at the forearm from the discharge timings decoded at the wrist. (**A**) Acquisition setup for concurrent recording of electromyogram (EMG) signals at the forearm and tendon electric signals at the wrist. (**B**) Representative EMG signal from a single contraction at the forearm. (**C**) Decoded motoneuron discharge timings from the tendon electric signals at the wrist for the same contraction. (**D**) Two representative examples of reconstructed motoneuron fiber action potentials for each channel at the forearm after spike trigger averaging the EMG signals across 50 ms windows centered at the discharge timings of the motoneurons detected at the wrist. The rationale for this approach is that if the discharge times decoded at the wrist correspond to the times of activation of spinal motoneurons, then the triggered average should identify muscle fiber potentials at the forearm above the baseline noise. The detection threshold was set to four times the baseline noise which is depicted for each channel as blue dotted lines. The channels that met this condition are framed in grey. As shown in the first example, only the channels that corresponded to muscle fiber action potentials were selected. Simultaneously, the detection condition was not met in the second example despite the variable amplitude levels, as no channel exhibited the stereotypical action potential waveform. This analysis showed that 703 out of 970 motoneurons decoded from the wrist were retraceable to the forearm, which proves the neural origin of the decoded tendon electric signals.

**Fig. S3.**
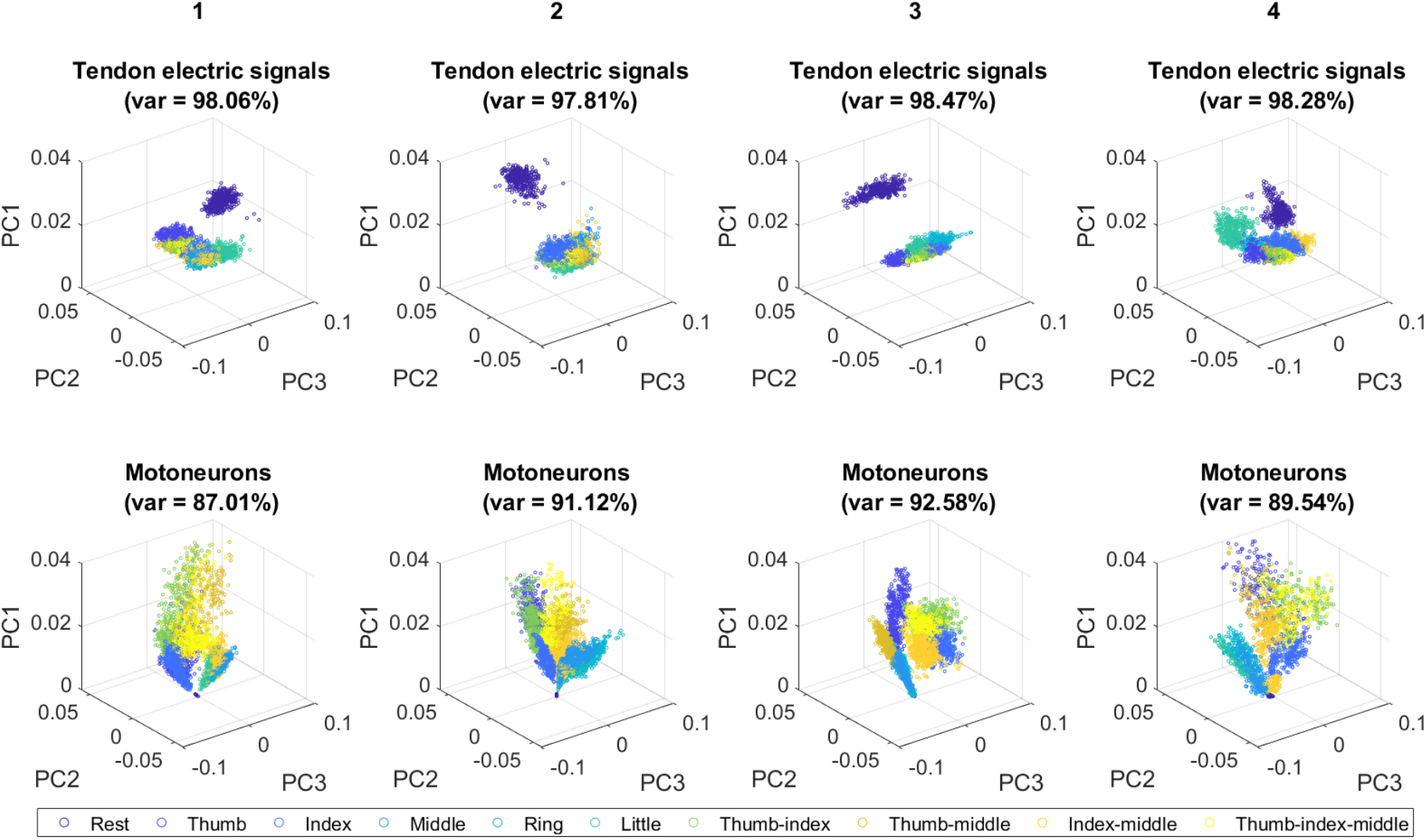
Feature space for the tendon electric signals and decoded motoneurons from the online task training. Visualization of the features for the tendon electric signals (top) comprising the root mean square, slope sign changes, waveform length, and zero crossings for each channel, and the spike count of the decoded motoneurons (bottom) over the first three principal components with the total explained variance between brackets. Each column represents one participant and finger contractions are color coded. The figure shows higher separability between finger contractions in the motoneuron feature space than in the tendon electric signals one.

