## Supplementary figures for "Non-invasive real-time access to the output of the spinal cord via a wrist wearable interface"

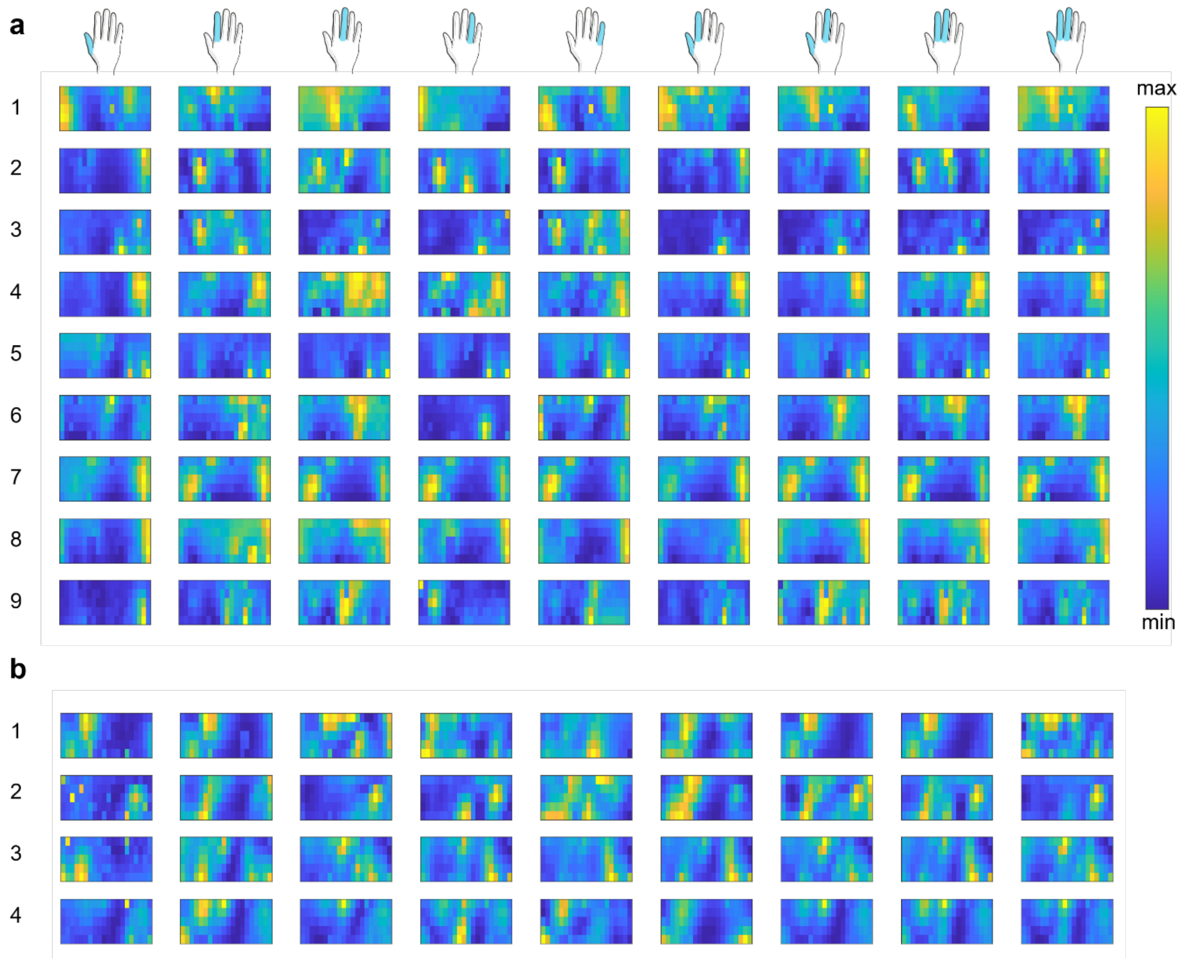


**Supplementary figure 1 | 2D spatial distribution of the normalised amplitude of the tendon electric signals.** Each subplot depicts the normalised root mean square of the tendon electric signals for each channel (pixel) in their corresponding spatial distribution in the electrode array at the wrist (view: bottom = proximal, top = distal, left = ulna posterior, right = ulna anterior). The values for few discarded channels due to noise have been estimated by 2D linear interpolation. The figure shows a high overlap in the activity area of the different finger contractions (columns) within each subject (rows). **a**, mean across the maps at 15% and 30% of force efforts from the offline dataset. **b**, mean across the maps of the three repetitions of the training set for the online prediction task.

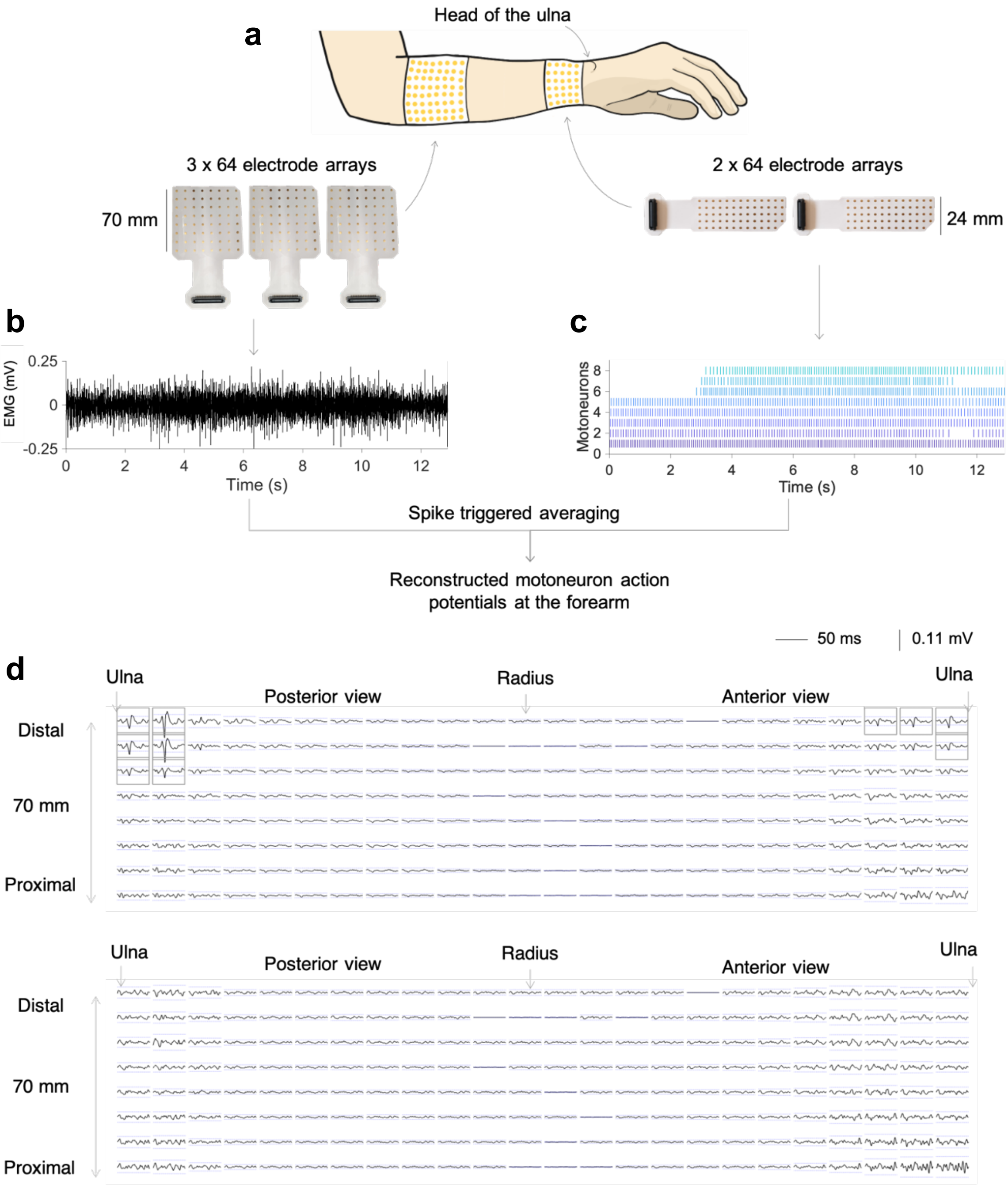


**Supplementary figure 2 | Retracing motoneuron fibre action potentials at the forearm from the discharge timings decoded at the wrist.** **a**, Acquisition setup for concurrent recording of electromyogram (EMG) signals at the forearm and tendon electric signals at the wrist. **b**, Representative EMG signal from a single contraction at the forearm. **c**, Decoded motoneuron discharge timings from the tendon electric signals at the wrist for the same contraction. **d**, Two representative examples of reconstructed motoneuron fibre action potentials for each channel at the forearm after spike trigger averaging the EMG signals across 50 ms windows centered at the discharge timings of the motoneurons detected at the wrist. The rationale for this approach is that if the discharge times decoded at the wrist correspond to the times of activation of spinal motoneurons, then the triggered average should identify muscle fibre potentials at the forearm above the baseline noise. The detection threshold was set to four times the baseline noise which is depicted for each channel as blue dotted lines. The channels that met this condition are framed in grey. As shown in the first example, only the channels that corresponded to muscle fibre action potentials were selected. Simultaneously, the detection condition was not met in the second example despite the variable amplitude levels, as no channel exhibited the stereotypical action potential waveform. This analysis showed that 703 out of 970 motoneurons decoded from the wrist were retraceable to the forearm, which proves the neural origin of the decoded tendon electric signals.

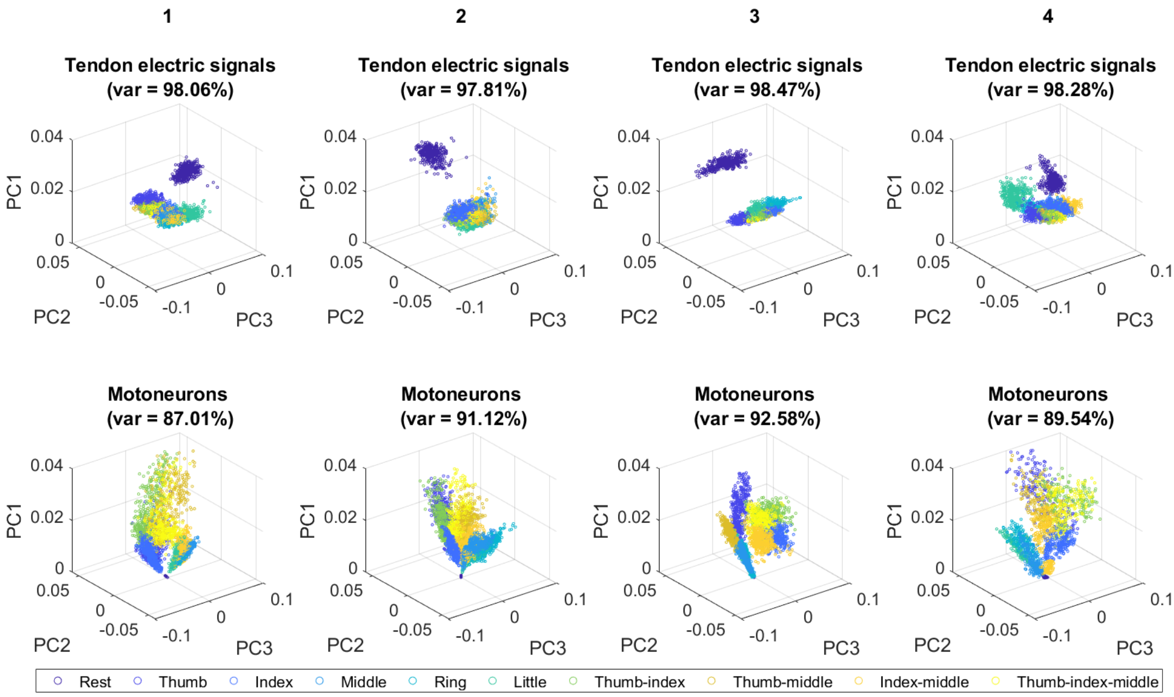


**Supplementary figure 3 | Feature space for the tendon electric signals and decoded motoneurons from the online task training.** Visualisation of the features for the tendon electric signals (top) comprising the root mean square, slope sign changes, waveform length, and zero crossings for each channel, and the spike count of the decoded motoneurons (bottom) over the first three principal components with the total explained variance between brackets. Each column represents one participant and finger contractions are colour coded. The figure shows higher separability between finger contractions in the motoneuron feature space than in the tendon electric signals one.
